# A curated dataset and web application for Integrative Analysis of Ancient DNA from Britain and Ireland

**DOI:** 10.1101/2025.05.30.657009

**Authors:** Sebastian Metz, Claire-Elise Fischer, Madeleine Bleasdale, Laura Castells Navarro, Lindsey Büster, Michael Legge, David Reich, Ian Armit

## Abstract

In recent years, genome-wide data from thousands of ancient human remains have been sequenced, offering unprecedented insights into past populations. This vast dataset of ancient human genomes and their associated metadata holds immense value for archaeologists, anthropologists, and biologists. However, extracting insights often requires advanced bioinformatics knowledge, creating barriers for non-experts. To address this challenge, we present a curated dataset and user-friendly web application for published genome-wide data from ancient Britain and Ireland, including new data produced by the COMMIOS Project. The platform integrates visualization tools, enabling researchers to explore genetic connections between ancient and modern populations without requiring bioinformatics expertise.

## Background & Summary

Over the past decade, the ‘Third Science Revolution’, centered around ancient DNA (aDNA) and big data, has transformed our understanding of the human past by providing invaluable insights into population evolution, migration and human-scale interactions^1,2^. Thousands of genome-wide samples are now accessible through public databases, making it an invaluable source of information for many fields of research, including archaeology, anthropology and biomedicine.

One of the major findings of this research has been that modern Europeans are a genetic mosaic of three major ancient populations: Western European hunter-gatherers (WHG), early European farmers (EEF) originating from Anatolia, and steppe pastoralists associated with the Yamnaya culture and their neighbours (Steppe)^3,4^, with much of the research published over the last decade revealing a complex series of expansions, replacements and admixture processes through time.

In a British context, aDNA has helped explore the impact of prehistoric and historic migrations. For example, the Beaker phenomenon at the beginning of the Bronze Age (c. 2,450 BCE) was associated with a major genetic transformation involving the movement of Bell Beaker culture-associated individuals into Britain. Between c. 2,400 - 2,000 BCE, these new communities, characterized by their Steppe ancestry, became numerically dominant to the extent that, by the Middle Bronze Age (1,800 - 800 BCE), Steppe ancestry contributed more than 90% of the genetic makeup of populations in Britain^5,6^. Following the Beaker period, during the Middle to Late Bronze Age, further substantial population movement into southern Britain from continental Europe occurred^7,8^. A recent study has also discussed the genetic impact of Viking migrations^9^. This work demonstrated that the Vikings were genetically highly diverse with significant gene flow from across Europe, challenging their traditional perception as a homogenous Scandinavian group^9^ and highlighting the role of Britain in the genetic composition of those groups. As a result of these and other research projects, thousands of ancient individuals from Britain and Ireland have now been sequenced^5,7–15^ (**Figure 1**).

**Figure 1.**
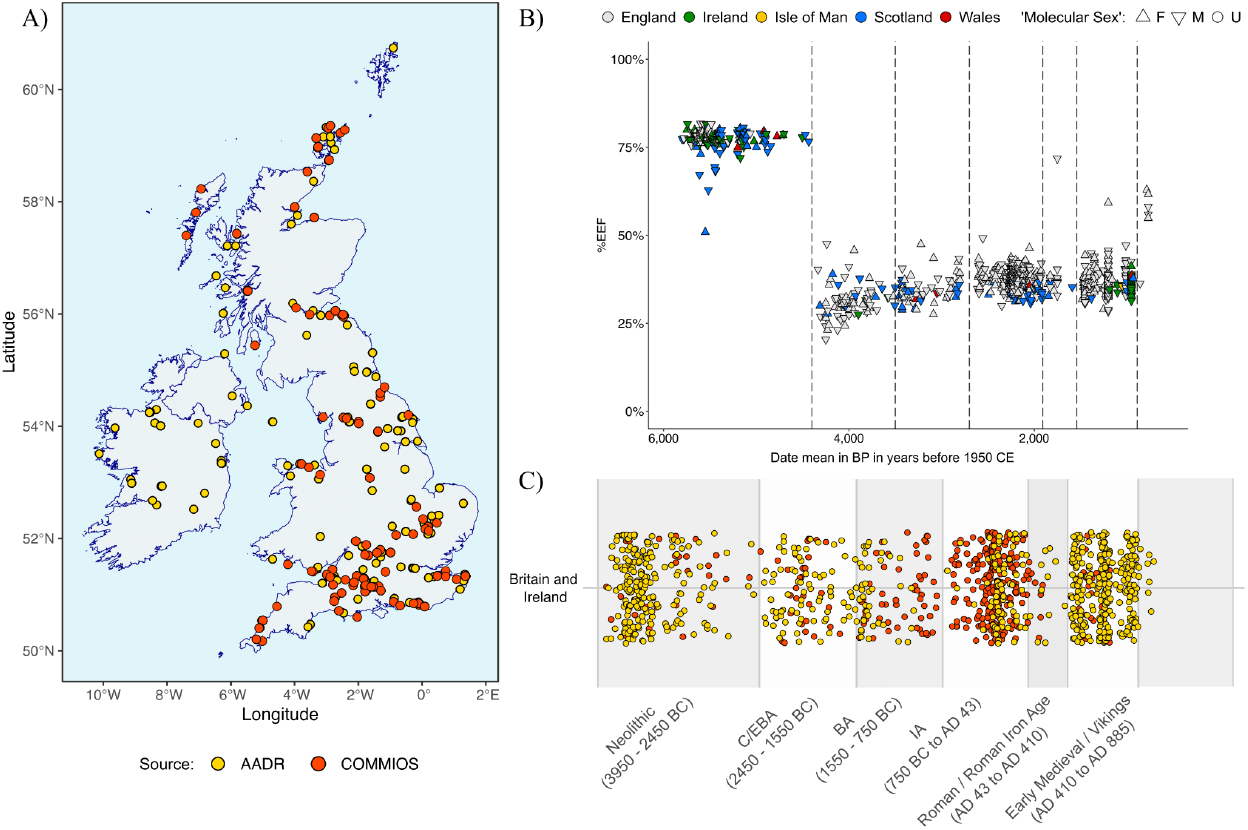
Overview of COMMIOS app dataset. a) Sampling distribution, including samples related to the COMMIOS Project (red) and those relating to other projects (yellow). b) Time series of the EEF contribution to ancient individuals from Britain and Ireland; patterning follows the results presented in Patterson et al.7 c) Sample distribution across different archaeological periods in Britain.

While the palaeogenetics community has made considerable efforts to ensure that databases are regularly updated and made publicly accessible (see for example AADR^16^ or Poseidon^17^), the use of these data by non-specialists is often restricted in practice. Accessing and analyzing aDNA data often requires advanced bioinformatics knowledge and tools which creates a barrier for researchers without expertise in this area. Furthermore, although research teams strive to provide the relevant meta-data (cultural period, radiocarbon date if available, age at death, genetic sex, etc.), it is generally archaeologists who have the expertise to fully understand the social context of the studied communities. In order to address some of these challenges, new platforms such as DORA^18^, MAPMIXTURE^19^ and ADGAP^20^ have been developed to facilitate the exploration of these complex datasets. However, to use most of these tools a certain level of technical expertise is still required.

Here we present the COMMIOS app, which draws on data collected by the COMMIOS Project (37.6% of the individuals) and individuals collected as part of other studies and collated in the AADR database (62.4%, **Figure 1**), and is a comprehensive, user-friendly tool that facilitates access to integrated data visualisation and exploration functionalities, lowering barriers for interpretation of the genetic dataset from Britain and Ireland.

## Methods

### Data Collection and Pre-processing

The dataset includes all ancient individuals from Britain and Ireland dating from 3,950 BCE to CE 1,200, for whom publicly available information exists^5,7,10–14,21–29^ (**Table 1**). Raw sequencing data are accessible via public repositories, in fastq or BAM file formats (**Table S1**). Genotypic data were extracted from the Allen Ancient DNA Resource v62 (AADR^16^), a comprehensive, curated database of ancient human genomes. All samples were processed using the AADR pipeline outlined in Mallick et al.^16^. This pipeline encompassed stringent quality control, alignment to the hg19 human genome reference, and genotype calling at approximately 1.23 million single nucleotide polymorphisms^3^ (SNPs).

**Table 1.** Britain and Ireland data sources, publications, number of samples and references.

| Original Publication | # of samples | Reference |
| --- | --- | --- |
| Brace *et al.*, 2019 | 36 | [10] |
| Cassidy *et al.*, 2020 | 42 | [11] |
| Olalde *et al.*, 2018 | 153 | [5] |
| Patterson *et al.*, 2022 | 412 | [7] |
| Fowler *et al.*, 2021 | 62 | [12] |
| Sánchez-Quinto *et al.*, 2019 | 15 | [21] |
| Scheib *et al.*, 2019 | 2 | [22] |
| Allentoft *et al.*, 2024 | 7 | [23] |
| Dulias *et al.*, 2022 | 37 | [13] |
| Cassidy *et al.*, 2016 | 4 | [24] |
| Rohland *et al.*, 2022 | 2 | [25] |
| Schiffels *et al.*, 2016 | 10 | [14] |
| Martiniano *et al.*, 2016 | 9 | [26] |
| Margaryan *et al.*, 2020 | 47 | [9] |
| Gretzinger *et al.*, 2022 | 251 | [28] |
| Brace *et al.*, 2022 | 6 | [29] |

### Metadata Curation and Temporal Classification

Each individual from the AADR underwent a manual review to identify and resolve inconsistencies in the metadata. The complete dataset used in the app includes 1,108 ancient British and Irish individuals (**Table 1**), 1,164 modern individuals and 4,618 ancient Western European individuals for purposes of comparison (**Table S1**). Temporal classification was performed based on radiocarbon dating or contextual archaeological evidence. Temporal periods for British and Irish individuals included in the application were defined as follows:

- **Neolithic**: 3,950–2,450 BCE
- **Chalcolithic/Early Bronze Age**: 2,450–1,550 BCE
- **Bronze Age**: 1,550–750 BCE
- **Iron Age (IA)**: 750 BCE – CE 43
- **Roman / Roman Iron Age**: CE 43 – 410
- **Viking/Early Medieval**: CE 410–1,100
- **Medieval**: CE 1,100–1,200

### Quality Assessment

Sample quality was evaluated for each individual based on the metadata available and the number of SNPs mapped to the 1240k SNPs and Human Origins (HO) SNP sets^3,30^. Each individual was then ascribed a value from 0 to 10 – based on individuals comprising more than 100k SNPs, mtDNA coverage, Molecular Sex identification and the “ASSESSMENT” classification from the AADR metadata. Individuals with high scores (> 8) were selected as references or good-quality samples, with the remainder flagged as “Quality check”, so as to alert the user of the app that these particular samples are of low quality or missing valuable information.

### Ancient proportions and Principal Component Analysis

Ancestry proportion estimates were obtained following Patterson et al.^7^ for all British and Irish ancient individuals, using the R package ADMIXTOOLS2^31^. Each individual was modelled as a mixture of Early European Farmers (EEF), Western Hunter-Gatherers (WHG), and Steppe pastoralists (Steppe). For this analysis, we use as the right group “OldAfrica”, “Afanasievo”, “Turkey_N” and “WHGB” and as the left group the three principal ancestry groups “OldSteppe”, “WHG” and “EEF”, following Patterson et al.^7^. In the case of Neolithic individuals we used a model of “WHG” and “EEF” only. Principal Component Analysis (PCA) was performed using smartPCA^32^. Selected modern individuals from the AADR were used as reference data, with ancient individuals projected onto the resulting PCA space^32^.

### App Development for Interactive Exploration

An interactive web application was developed using Shiny R^33^ to facilitate the exploration and visualization of the results. The app provides the following key functionalities:

- **Sample Distribution Map**: A geospatial visualization displaying the distribution of ancient samples across regions (**Figure 1a**).
- **Temporal Analysis of Admixtures**: Visualizations of temporal trends in population contributions and shifts in ancestry patterns based on admixture models for each individual, showing the contributions from major ancient populations such as Early European Farmers (EEF), Western Hunter-Gatherers (WHG) and Steppe pastoralists (**Figure 1b**).
- **Metadata Table**: A searchable table containing manually curated metadata for all samples, including temporal, geographic, and quality assessments. These data are also available to access in GitHub / Zenodo in a table-separated format with the associated genotypes in EIGENSTRAT format^3,16^.
- **Principal Component Analysis (PCA)**: Interactive PCA plots to select reference populations and visualize ancient British samples in comparison to modern or ancient populations from Europe (**Figure S1**).
- **Ternary Plot**: A ternary plot visualizing the proportional contributions of ancient populations (EEF, WHG and Steppe) for all samples.

### Computational and Statistical Analyses

All data preprocessing, statistical analyses and visualizations were performed using R (version 4.4.0). Custom scripts and reproducible workflows are available in a GitHub repository (https://github.com/sebametz/COMMIOS_app).

## Data Records

The dataset and associated metadata are available through public repositories. The data include:

- **Curated Metadata** (**Table S1**): A manually curated dataset with detailed information on geographic location, temporal classification, sample quality, raw sequencing data accession numbers, and other relevant attributes.
- **Genotypes**: Provided in EIGENSTRAT format, including genotype calls for approximately 1.23 million SNPs and the Human Origins array with 597,573 sites. These datasets can be downloaded from Zenodo (10.5281/zenodo.15536132).
- **Analysis outputs**: Admixture proportions, PCA coordinates, and ternary plot values for each sample can be downloaded from the app in SVG format.
- **Scripts**: pre-processing R script, reference files, and the application script can be accessed from the public GitHub repository (https://github.com/sebametz/COMMIOS_app).
- **Web application**: The web application can be accessed from the University of York Shiny R services at https://shiny.york.ac.uk/COMMIOS_app/

## Technical Validation

Here, we presented a publicly available genome-wide dataset processed using the same pipeline for consistency and reproducibility. The metadata of each individual was manually curated and all changes can be tracked in the pre-processing scripts. Stringent filtering was performed to detect samples with insufficient SNP coverage and ambiguous metadata. A manual review of location and temporal classification was performed to ensure accuracy. Comparison of the Principal Component Analysis and admixture modelling results against published studies confirms consistency with prior findings^8^ (**Figure 1** and **Figure S1**).

The Shiny R application facilitates further validation by enabling interactive exploration and visualization of the data.

## Usage Notes

The dataset and accompanying app are designed to support a wide range of research questions in ancient DNA and population genetics. Potential applications include:

- Population Structure Analysis: Using PCA and admixture models to study genetic relationships among ancient populations.
- Temporal Trends: Examining changes in genetic contributions over time within and between regions.
- Geospatial Analysis: Investigating the geographic distribution of genetic variation using the sample distribution map.
- Customized Subset Analyses: Filtering samples by temporal period, geographic location or quality to focus on specific research questions.

Researchers are encouraged to use the Shiny R app for exploratory data analysis and hypothesis generation.

## Code Availability

The codes for data collection pre-processing and from the application are freely accessible on the GitHub repositories at https://github.com/sebametz/COMMIOS_app_preprocessing and https://github.com/sebametz/COMMIOS_app. The application can be accessed at https://shiny.york.ac.uk/COMMIOS_app/.

## Acknowledgements

Research for this article received funding from the European Research Council (ERC), under the European Union’s Horizon 2020 research and innovation programme, grant agreement no. 834087 (COMMIOS). The authors would like to thank Nathan Wales and Eleanor Green for their comments and feedback on the application functionality.

## Author contributions

S.M. designed and programmed the app; S.M. and I.A. wrote the paper with input from all authors.

---

**Figure S1.**
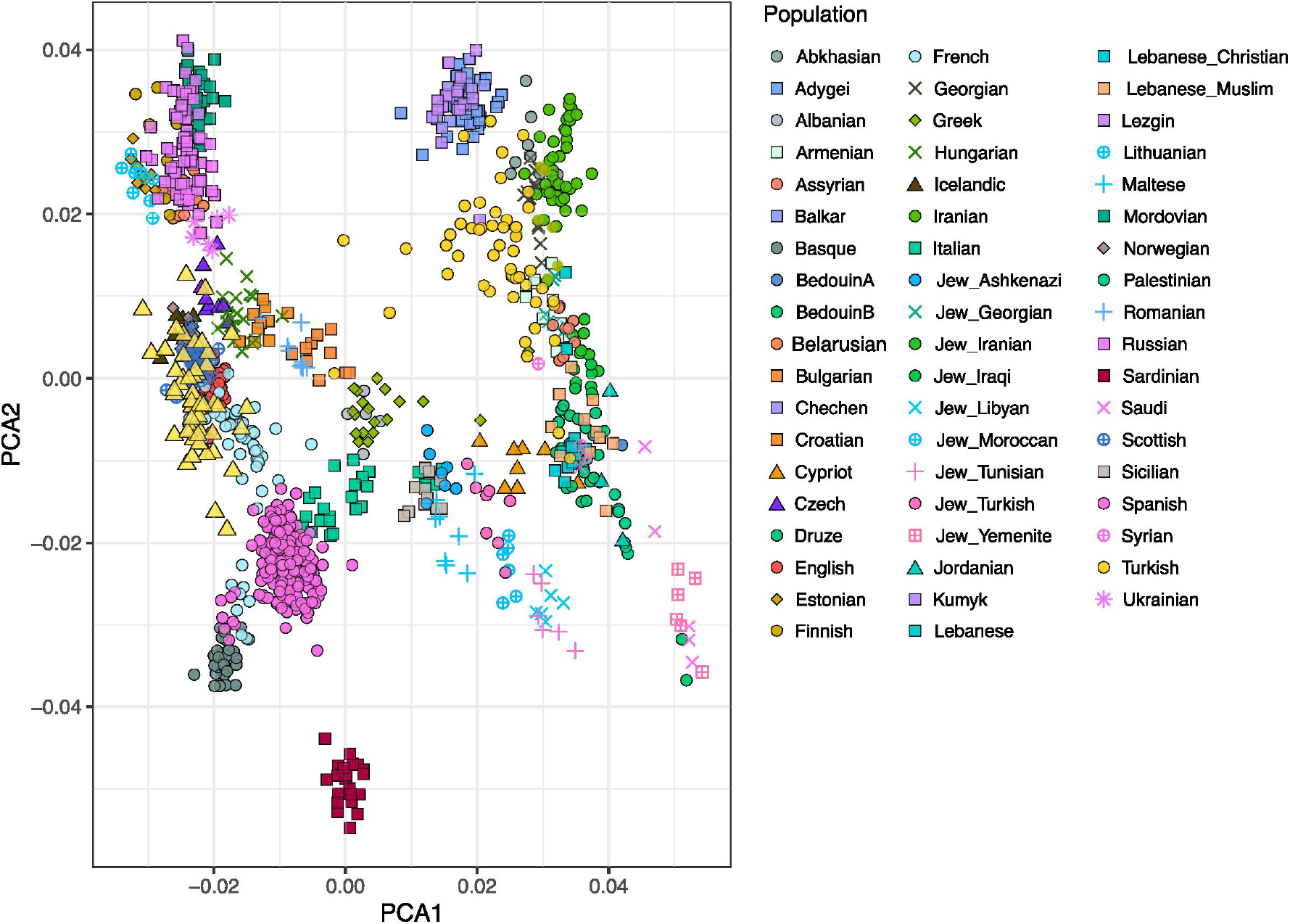

