## Supplementary material for "A curated dataset and web application for Integrative Analysis of Ancient DNA from Britain and Ireland": Figure S1

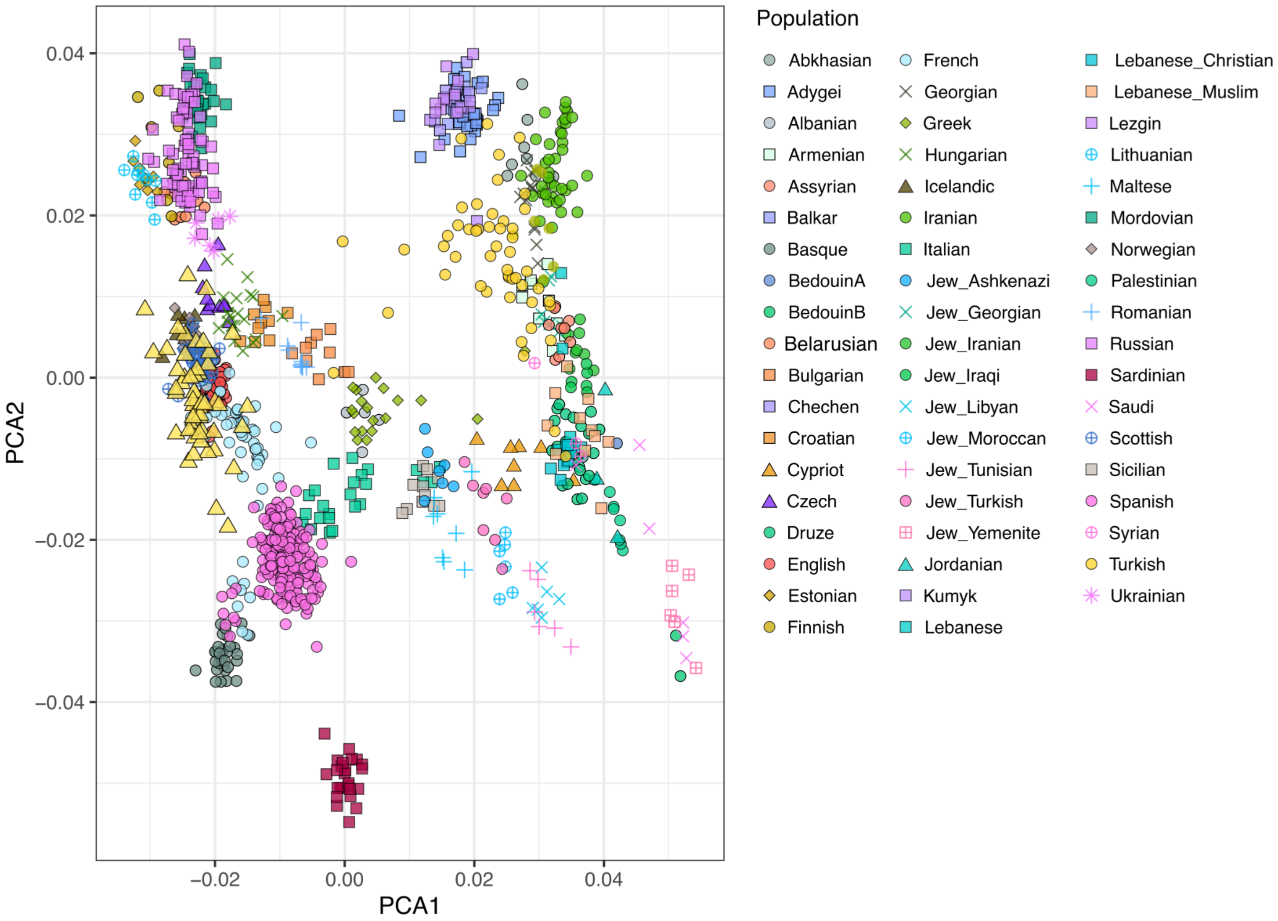


**Figure S1**: PCA distribution of modern individuals aligns consistently with the structures previously described by Lazaridis et al.[30]. Individuals from Kent, England (yellow triangles) exhibit a distribution consistent with expectations documented in prior studies.
